# Restriction and recruitment – gene duplication and the origin and evolution of snake venom toxins

**DOI:** 10.1101/006023

**Authors:** Adam D Hargreaves, Martin T Swain, Matthew J Hegarty, Darren W Logan, John F Mulley

**Author notes:** Corresponding author: Dr John Mulley, School of Biological Sciences, Bangor University, Deiniol Road, Bangor, Gwynedd LL57 2UW, United Kingdom.

## Abstract

The genetic and genomic mechanisms underlying evolutionary innovations are of fundamental importance to our understanding of animal evolution. Snake venom represents one such innovation and has been hypothesised to have originated and diversified via a process that involves duplication of genes encoding body proteins and subsequent recruitment of the copy to the venom gland where natural selection can act to develop or increase toxicity. However, gene duplication is known to be a rare event in vertebrate genomes and the recruitment of duplicated genes to a novel expression domain (neofunctionalisation) is an even rarer process that requires the evolution of novel combinations of transcription factor binding sites in upstream regulatory regions. This hypothesis concerning the evolution of snake venom is therefore very unlikely. Nonetheless, it is often assumed to be established fact and this has hampered research into the true origins of snake venom toxins. We have generated transcriptomic data for a diversity of body tissues and salivary and venom glands from venomous and non-venomous reptiles, which has allowed us to critically evaluate this hypothesis. Our comparative transcriptomic analysis of venom and salivary glands and body tissues in five species of reptile reveals that snake venom does not evolve via the hypothesised process of duplication and recruitment of body proteins. Indeed, our results show that many proposed venom toxins are in fact expressed in a wide variety of body tissues, including the salivary gland of non-venomous reptiles and have therefore been restricted to the venom gland following duplication, not recruited. Thus snake venom evolves via the duplication and subfunctionalisation of genes encoding existing salivary proteins. These results highlight the danger of the “just-so story” in evolutionary biology, where an elegant and intuitive idea is repeated so often that it assumes the mantle of established fact, to the detriment of the field as a whole.

## Introduction

Gene duplication is a rare event in eukaryotic genomes and has been suggested to be the major source of novel genetic material (Ohno 1970). Estimates of the rate of gene duplication in vertebrates vary from 1 gene per 100 to 1 gene per 1000 per million years (Lynch and Conery 2000; Lynch and Conery 2003; Cotton and Page 2005), and the most common fate for a duplicate gene is the loss of its function (nonfunctionalisation, pseudogenisation (Mighell et al. 2000; Presgraves 2005)). However, in some cases a duplicate gene is retained in the population and undergoes either subfunctionalisation (where the two duplicates divide the sum of the ancestral role(s) between them) or neofunctionalisation (where one of the duplicates assumes a new role, independent of the ancestral function (Force et al. 1999)). This latter process of evolving an entirely new function is known to be incredibly rare and there are few conclusive examples of it in the literature (Escriva et al. 2006; Van Damme et al. 2007; Deng et al. 2010).

The venom of advanced snakes has been hypothesised to have originated and diversified via gene duplication (Wong and Belov 2012). In particular, it has been suggested that both the origin of venom and the later evolution of novelty in venom has occurred as a result of the duplication of a gene encoding a non-venom physiological or “body” protein that is subsequently recruited, via gene regulatory changes, into the venom gland, where natural selection can act on randomly occurring mutations to develop and/or increase toxicity (Lynch 2007; Fry et al. 2009b; Kwong et al. 2009; Casewell et al. 2012; Fry et al. 2012a; Casewell et al. 2013; Margres et al. 2013; Vonk et al. 2013). In short, it has been proposed that snake venom diversifies via repeated gene duplication and neofunctionalisation, a somewhat surprising finding given the apparent rarity of both of these events (here we refer to neofunctionalisation with respect to the acquisition of novel sites of expression at the level of individual tissues, not the acquisition of novel functions at a molecular level, which is separate from the claims of the duplication/recruitment hypothesis and has been shown to have occurred for only a small number of venom toxins (Kini 2002; Kini 2003; Lynch 2007; Kini and Doley 2010), whilst the majority of duplicated toxins retain ancestral bioactivity (Fry 2005; Warrell 2010)). However, there are currently several gaps in our knowledge of how this remarkable process might take place, including the mechanisms underlying repeated gene duplications and, more importantly, the gene regulatory changes that occur to facilitate “recruitment” into the venom gland. Given that whole genome duplication is a rare event in vertebrates in general and reptiles in particular (Otto and Whitton 2000; Mable 2004), it seems likely that the majority of snake venom toxin genes are duplicated via segmental duplication (Hurles 2004), where the highly repetitive nature of reptile genomes (Shedlock et al. 2007; Di-Poi et al. 2009) provides regions of pseudo-homology that facilitate unequal crossing-over during homologous recombination, producing tandemly-arranged duplicates. This process requires neither germ-line expression nor the evolution of *de novo cis*-regulatory sequences as does retrotransposition (Zhang 2003) and, if repeated so that the resulting pairs or larger clusters of genes were subsequently duplicated in the same manner, a relatively small number of duplication events could give rise to a large number of duplicate genes. Evidence for clusters of multiple SVMP, CRISP and lectin genes in the king cobra genome (Vonk et al. 2013) and for PLA_2_ genes in the Okinawan habu (*Protobothrops* (now *Trimeresurus*) *flavoviridis*) (Ikeda et al. 2010) would seem to support this hypothesis, although more complete data from these and other snake whole genome sequencing projects is needed.

Whilst the above scenario explains the apparent ease with which *existing* venom toxin genes might be repeatedly duplicated along with their associated *cis*-regulatory architecture, it does nothing to explain how a non-venom gene might be “recruited” into the venom gland. The paralogous genes produced as a result of gene duplication are 100% identical and, if the entirety of their associated *cis*-regulatory architecture has also been duplicated along with them, they will have identical temporal and spatial expression patterns (i.e. they are functionally redundant (Force et al. 1999; Lynch and Force 2000)). Therefore in order to develop a novel site of expression such as in the venom gland, a novel combination of transcriptional regulatory sequences must arise.

Eukaryotic transcription factor binding sites are the result of a trade-off between the specificity offered by longer stretches of DNA and the robustness to mutation offered by shorter sequences and vary in length between 5 and >30nt, with an average length of 10nt (Stewart et al. 2012). It has been estimated that eukaryotic promoters may contain 10-50 binding sites for 5-15 different transcription factors (Wray et al. 2003). The rarity of gene duplication, coupled with the low likelihood of evolving new combinations of transcription factor binding sites before the duplicated gene is nonfunctionalised by random mutations in coding sequences should therefore make the process of duplication and recruitment of genes encoding physiological or body proteins into the venom gland exceedingly rare. How then do we reconcile this with the apparent widespread occurrence of just this process in the origin and evolution of snake venom? One possible alternative hypothesis is that many of the genes expressed in snake venom are in fact the result of the duplication of genes that were ancestrally expressed in multiple tissues, including the venom gland. Following duplication these genes therefore evolved via subfunctionalisation, with one copy’s expression being restricted to the venom gland and the other maintaining the original, multi-tissue expression pattern (possibly with subsequent loss of expression of this paralog in the venom gland). This scenario of duplication and restriction, rather than duplication and recruitment (Figure 1) is more parsimonious as it requires only the loss of transcription factor binding sites, which may occur by random mutation of single base pairs or larger insertions or deletions (indels) that may delete or disrupt the existing transcriptional regulatory sequences. In order to differentiate between the two hypotheses gene expression data from non-venom gland tissues in venomous and non-venomous species are needed, something which has until now been missing. Here we review the existing evidence for the duplication and recruitment of genes into the venom gland and carry out a comparative transcriptomic survey of gene expression in the venom glands and body tissues of a number of reptile species, including the painted saw-scaled viper (*Echis coloratus*), a venomous snake; the corn snake (*Pantherophis guttatus*) and rough green snake (*Opheodrys aestivus*), both non-venomous colubrids which use constriction to kill prey (Kardong 2002); the royal python (*Python regius*) a non-venomous boid and the leopard gecko (*Eublepharis macularius*), a member of one of the most basal lineages of squamate reptiles. The phylogenetic position of this latter species is particularly important, as it lies outside of the proposed clade of ancestrally venomous reptiles (the Toxicofera (Fry et al. 2006; Fry et al. 2009a; Fry et al. 2012b; Fry et al. 2013)) and therefore genes found in the salivary gland of this species can be taken to represent the ancestral squamate expression pattern. We also take advantage of available transcriptomic resources for body tissues in a number of other reptile species, including king cobra (*Ophiophagus hannah*) venom gland, accessory gland and pooled tissues (heart, lung, spleen, brain, testes, gall bladder, pancreas, small intestine, kidney, liver, eye, tongue and stomach) (Vonk et al. 2013), garter snake (*Thamnophis elegans*) liver (Schwartz and Bronikowski 2013) and pooled tissue (brain, gonads, heart, kidney, liver, spleen and blood of males and females) (Schwartz et al. 2010), Burmese python (*Python molurus bivittatus*) pooled heart and liver (Castoe et al. 2011) and corn snake brain (Tzika et al. 2011).

**Figure 1.**
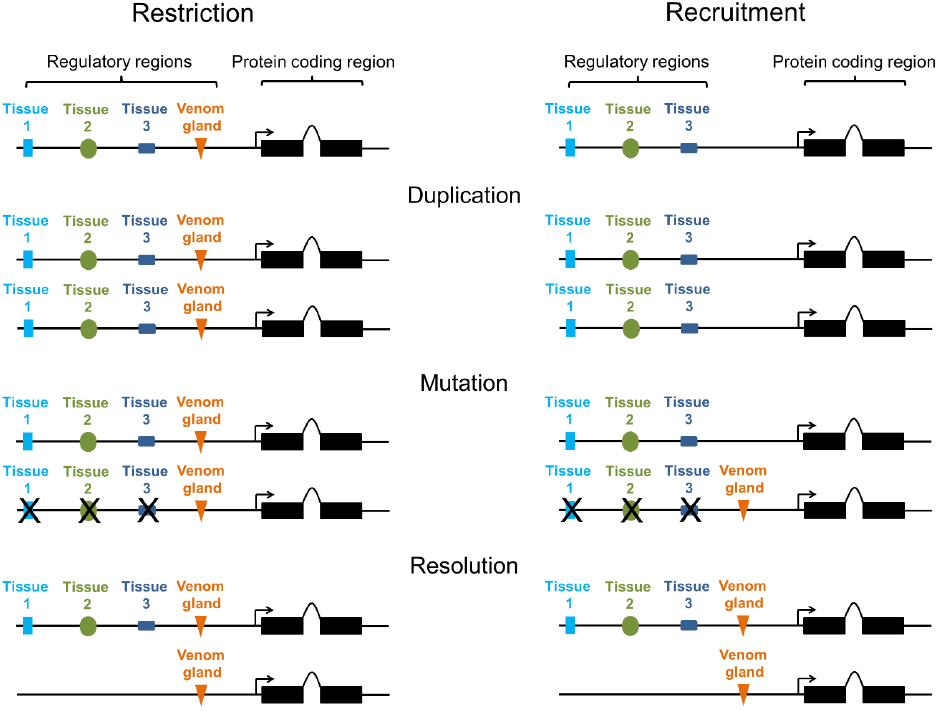
Restriction and recruitment. Duplicated genes may be either restricted or recruited to the venom gland, with the latter process dependent on the evolution of new combinations of transcription factor binding sites in upstream regulatory regions. Mutation/loss of regulatory regions is indicated with an X.

## Results and Discussion

We find the hypothesis that snake venom evolves via the duplication of physiological or body genes and subsequent recruitment into the venom gland to be unsupported by the available data – in short, snake venom has not evolved via the recruitment of “body” genes. Indeed for a large number of the gene families claimed to have undergone recruitment we find evidence of a diverse tissue expression pattern, including the salivary gland of non-venomous reptiles (Figure 2), demonstrating that, if they do encode toxic venom components (Hargreaves *et al*. in prep), they have not been recruited into the venom gland, but restricted to it. The recently published king cobra genome paper (Vonk et al. 2013) also provides evidence for salivary (rictal) gland expression of several venom toxins in the Burmese python, *Python molurus bivittatus*, including 3ftx, cystatin, hyaluronidase and SVMP (Supplementary Table S2 in (Vonk et al. 2013)).

**Figure 2.**
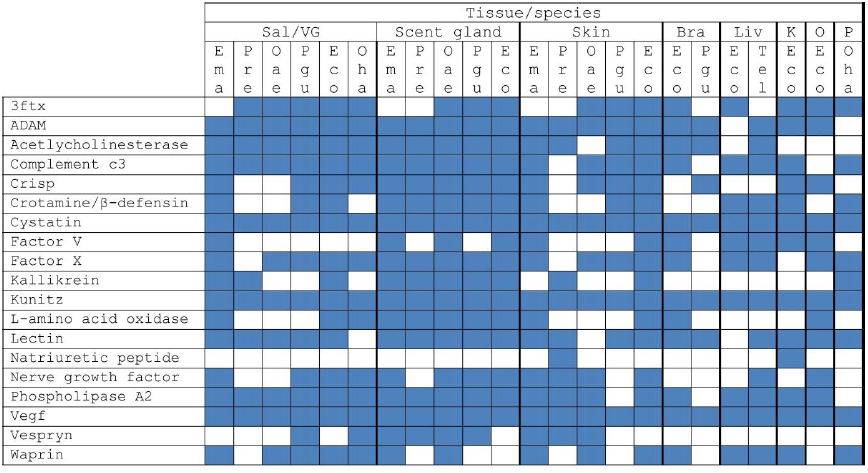
Tissue distribution of putative toxin gene families. Many proposed toxin gene families are expressed in a wide range of tissues, including the salivary or venom gland and have therefore been restricted to the venom gland following duplication, not recruited. Tissue abbreviations: Sal, salivary gland; VG, venom gland; Bra, brain; Liv, liver; K, kidney; O, ovary; P, pooled tissue (see text for details). Species abbreviations: Ema, leopard gecko (*Eublepharis macularius*); Pre, royal python (*Python regius*); Oae, rough green snake (*Opheodrys aestivus*); Pgu, corn snake (*Pantherophis guttatus*); Eco, painted saw-scaled viper (*Echis coloratus*); Oha, king cobra (*Ophiophagus hannah*); Tel, garter snake (*Thamnophis elegans*).

Therefore whilst some venom toxin genes have in the past been suggested to represent ancestral salivary proteins (notably cysteine-rich secretory proteins (CRISPs)) and Kallikrein-like serine proteases (Fry 2005; Sunagar et al. 2012), our analysis in fact shows that the majority of snake venom toxins are likely derived from pre-existing salivary proteins. Far from being an incredibly complex cocktail of proteins (Kini 2002; Wagstaff et al. 2006; Fox and Serrano 2008; Casewell et al. 2013) recruited from multiple body tissues (Fry 2005; Fry et al. 2009a; Warrell 2010; Casewell et al. 2013), snake venom should instead be considered to be simply a modified form of saliva, where a relatively small number of gene families (typically 6-14) have expanded via gene duplication, often in a lineage-specific manner (Kulkeaw et al. 2007; Wagstaff et al. 2009; Fahmi et al. 2012; Vonk et al. 2013).

The study cited most frequently in support of the duplication and recruitment hypothesis is that of Fry (Fry 2005) (see for example (Warrell 2010; Jiang et al. 2011; Casewell et al. 2012; Casewell et al. 2013)) and we therefore refer to this hypothesis as the ‘genome to venome hypothesis’. In his study, Fry concluded that the evolution of snake venom was characterised by at least 24 recruitment events (Fry 2005). However, this analysis was based on assumptions that snake venom toxin sequences derived primarily from EST-based studies of only the venom gland could be considered to be venom gland-specific and that if they were related to a gene known to be expressed in the pancreas (or another tissue) of human or other species they must therefore represent a recruitment event. It is obviously possible that the same gene may be expressed in the pancreas (or other tissue) of the snake as well and that the lack of data for these non-venom gland tissues is obscuring the true extent of their expression. It must be considered therefore that for the majority of genes Fry does not actually demonstrate any evidence for gene duplication and subsequent recruitment.

Only four examples in Fry’s study include both “body” and venom gland sequences from venomous snakes and therefore only these four possibly show any evidence in support of gene duplication and recruitment into the venom gland: crotamine; complement C3; natriuretic peptide and Group IB phospholipase A_2_ (Fry 2005). Of these, the South American rattlesnake (*Crotalus durissus terrificus*) *crotamine-*like sequence labelled as ‘Pancreas’ (accession number Q6HAA2) was in fact originally described to be highly expressed in pancreas, heart, liver, brain and kidneys (i.e. all tissues examined) with “scarce” but detectable expression in the venom gland (Rádis-Baptista et al. 2004). Our transcriptomic data shows that the toxic form of *crotamine* is derived from the duplication of a non-toxic *β-defensin*-like gene with a wider expression pattern that included the salivary/venom gland (Figure 2) and that the toxic duplicate has been restricted, not recruited, to the venom gland. For *complement C3*, Fry’s analysis utilised Indian cobra (*Naja naja*) sequences from liver (accession number Q01833) (Fritzinger et al. 1992) and venom gland (accession number Q91132) (Fritzinger et al. 1994). However, both sequences were in fact isolated from what the authors refer to as “*Naja naja kaouthia*”, a synonym for the monocled cobra, *Naja kaouthia*. This inaccuracy notwithstanding, Fry’s analysis does suggest that there has been a duplication of a *complement C3* gene to give rise to a new copy (often referred to as “*cobra venom factor*”, more rightly called *complement C3b*) although the lack of data for other body tissues should have precluded claims of recruitment. Analysis of our transcriptome data in fact reveals that *complement C3* is expressed in a diverse array of body tissues in multiple species, including the salivary gland of non-venomous reptiles (Figures 2 and 3) and that a paralogous copy of this gene has therefore been restricted to the venom gland following duplication. Whilst *Bothrops jararaca* does appear to possess at least two distinct forms of natriuretic peptide (Hayashi et al. 2003; Hayashi and Camargo 2005), the situation may also be more complex than that originally presented, as the sequence labelled as ‘Brain’ by Fry (accession Q9PW56, identical to AAD51326) in fact shows a wider expression pattern that includes brain, spleen, venom gland and, possibly, pancreas (Murayama et al. 1997; Hayashi et al. 2003; Hayashi and Camargo 2005). We find few natriuretic peptides in our dataset (Figure 2), and the low number of these sequences previously characterised would suggest that they play little role in the venom of non-*Bothrops* snakes, where they appear to have undergone duplication and subfunctionalisation. Finally, Fry used *Group IB phospholipase A_2_* (*PLA_2_ IB*) sequences from the pancreas of the banded sea krait (*Laticauda semifasciata*, accession Q8JFG2) and the venom gland of the Australian coastal taipan (*Oxyuranus scutellatus*, accession P00615) to support recruitment. We find *PLA_2_ IB* genes to be expressed in several body tissues, including the leopard gecko salivary gland (Figure 2 and Supplementary figure 1), suggesting a wider ancestral expression pattern than previously claimed.

**Figure 3.**
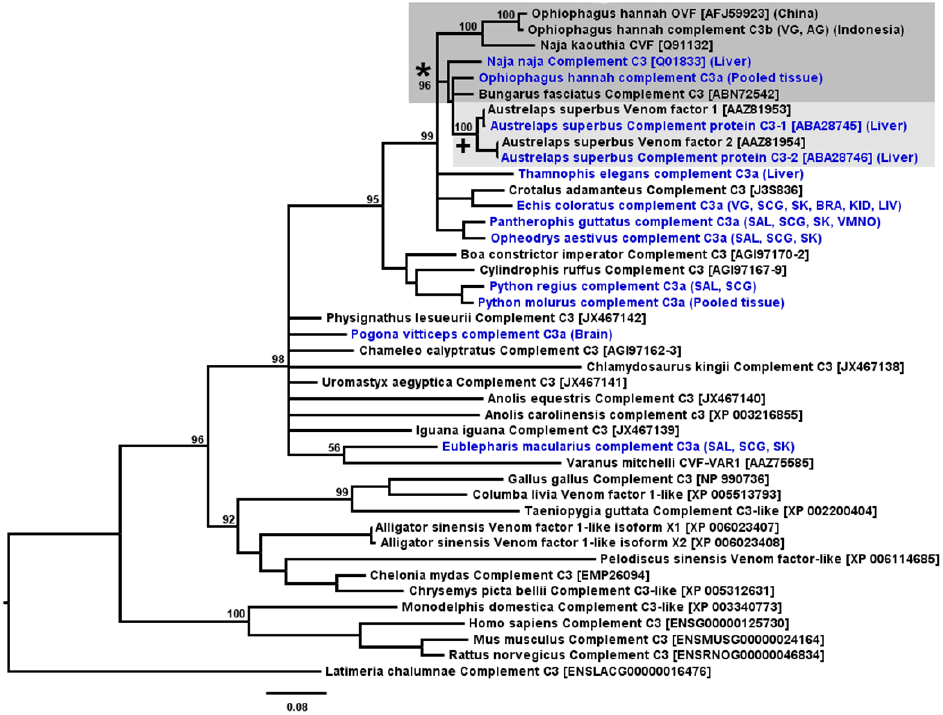
Maximum likelihood tree of *complement C3* genes. *complement C3* genes are expressed in a diversity of tissues, including venom and salivary glands. Following a gene duplication event (marked with *, shaded dark grey) one paralog has been restricted to the venom gland in the king cobra (*Ophiophagus hannah*) and the monocled cobra (*Naja kaouthia*). The two distinct king cobra sequences most likely represent geographic variation between Indonesian and Chinese populations. An additional gene duplication event appears to have occurred in the *Austrelaps superbus* lineage (marked with +, shaded light grey). Lineages for which body (non-venom gland) sequences are available are coloured blue and bootstrap values for 500 replicates are shown above branches.

It has recently been suggested that there has been a duplication of *nerve growth factor* (*ngf*) genes in some species of snake (Sunagar et al. 2013), although the presence of additional copies of *ngf* in certain species of cobra has been known for some time (Lipps 2000; Koh et al. 2004). Our data show that the non-toxic form of *ngf* (which we call *ngfa*) is expressed in a diversity of tissues, including the salivary glands of non-venomous reptiles (Figure 2 and Supplementary figure 2). The putatively toxic version (*ngfb*) has therefore also been restricted to the venom gland following duplication.

Both coagulation *factor V* and *factor X* have been suggested to have undergone gene duplication in Australian elapids such as *Tropidechis carinatus* and *Pseudonaja textilis* with subsequent recruitment of a gene normally expressed in the liver into the venom gland (Le et al. 2005; Reza et al. 2007; Kwong et al. 2009; Kwong and Kini 2011). However, these studies do not appear to have investigated body tissues other than liver and venom gland (Le et al. 2005) and so cannot be relied upon to demonstrate the full extent of ancestral gene expression. Our analysis in fact shows *factor V* to be expressed in multiple tissues, including rough green snake scent gland, King cobra accessory gland, *Echis coloratus* scent gland, kidney, brain, ovary and skin and the scent gland, skin and salivary gland of the leopard gecko (Figure 2 and Supplementary figure 3). *Factor X* is also expressed in multiple tissues (Figure 2 and Supplementary figure 4), including the salivary or venom glands of leopard gecko, royal python, rough green snake, corn snake and *Echis coloratus*. In both cases therefore a gene with a wide expression pattern that included the salivary or venom gland has undergone duplication and restriction. The known increased expression of a *factor X* paralog following an insertion in the promoter region (Reza et al. 2007; Kwong and Kini; Kwong et al. 2009; Han et al. 2013) and the increased expression of *crotamine* in the venom gland following duplication (Rádis-Baptista et al. 2003; Rádis-Baptista et al. 2004) suggest that a possible route for pre-existing salivary proteins to become venom toxins may simply be an elevated expression level, where initial toxicity is dosage-dependent.

Interestingly, some of the key papers cited in support of the genome to venome hypothesis in fact discuss the recruitment of genes into the venom proteome, *not* the venom gland itself (Fry and Wuster 2004; Fry 2005) with such claims only becoming more common in the literature some time later (see for example (Fry et al. 2008; Durban et al. 2011; Casewell et al. 2013)). Added to the fact that these papers show no evidence for duplication and recruitment of “body” genes it must be concluded that not only is this hypothesis not supported by our newly available data, but that it was never supported. It appears therefore that a misunderstanding of the scope of the claims of these earlier studies, together with the known role for gene duplication in the *diversification* of snake venom (Kordiš and Gubenšek 2000) is responsible for the development and propagation of the attractive, but ultimately unsupported, duplication and venom gland recruitment hypothesis. In order to fully understand the evolution of snake venom, more transcriptomic data is needed from a much greater variety of species for a much greater number of body tissues, ideally at a wider diversity of stages of venom synthesis and with consideration of sex, ontogeny, shedding and reproductive cycles and the large-scale effects on metabolism of intermittent feeding on large prey (Wall et al. 2011; Castoe et al. 2013). Even so, it will be difficult to fully account for all possible spatial and temporal influences on gene expression, and the default assumption for the fate of duplicate genes should perhaps therefore be subfunctionalisation, not neofunctionalisation.

Finally, our findings highlight the problem of ‘just-so stories’ (Kipling 1902) in evolutionary biology, especially when they reach the point of being considered established fact. The genome to venome hypothesis has been widely and unquestioningly cited and treated neither as a hypothesis to be tested and refuted (Popper 1959), nor as a scientific research programme to provide predictions to be investigated (Lakatos 1980). Whilst the role of gene duplication should rightly be considered as part of the core of the snake venom evolution research programme, we propose that many associated hypotheses are in need of a greater degree of scrutiny than they have hitherto received. Only after such scrutiny will we truly understand “How The Snake Got His Venom”.

## Materials and Methods

Total RNA was extracted from the salivary glands, scent glands and skin of two adult corn snakes (*Pantherophis guttatus*), rough green snakes (*Opheodrys aestivus*), royal pythons (*Python regius*) and leopard geckos (*Eublepharis macularius*). Only a single corn snake skin sample provided RNA of high enough quality for sequencing. RNA samples for painted saw-scaled vipers (*Echis coloratus*) were extracted from the skin, scent glands, kidney and brain of two adult specimens, and liver and ovary samples were extracted from one adult individual. Venom glands from four adult individuals were taken at different time points following venom extraction (16, 24 and 48 hours post-milking) in order to capture the full diversity of venom genes. All RNA extractions were carried out using the RNeasy mini kit (Qiagen) with on-column DNase digestion. mRNA was prepared for sequencing using the TruSeq RNA sample preparation kit (Illumina) with a selected fragment size of 200-500bp and sequenced using 100bp paired-end reads on the Illumina HiSeq2000 or HiSeq2500 platform. The quality of all raw sequence data was assessed using FastQC (Andrews 2010) and reads for each tissue pooled and assembled using Trinity (Grabherr et al. 2011) (sequence and assembly metrics are provided in Supplementary tables S1 and S2). Venom genes were identified by BLAST (Camacho et al. 2009) and maximum-likelihood-based phylogenetic analysis and tissue distribution identified by BLAST-based searches of assembled transcriptomes.

Transcriptome reads were deposited in the European Nucleotide Archive (ENA) database under accession #ERP001222 and the Sequence Read Archive (SRA) under the study accession #SRP042007.

## Acknowledgements

The authors wish to thank R. Morgan, A. Barlow and C. Wüster for technical assistance and S. Webster and W. Wüster for comments on the manuscript. We would also like to acknowledge the always enthusiastic help and support of the late Ashley Tweedale. We are very grateful to the staff of High Performance Computing (HPC) Wales for enabling and supporting our access to their systems. This work was partially supported by a Royal Society Research Grant awarded to JFM (grant number RG100514) and Wellcome Trust funding to DWL (grant number 098051). JFM, MJH and MTS are supported by the Biosciences, Environment and Agriculture Alliance (BEAA) between Bangor University and Aberystwyth University and ADH is funded by a Bangor University 125^th^ Anniversary Studentship.

